# *Pseudomonas aeruginosa* senses and responds to epithelial potassium flux via Kdp operon to promote biofilm biogenesis

**DOI:** 10.1101/2023.06.05.543669

**Authors:** Glenn J Rapsinski, Madison Hill, Kaitlin D Yarrington, Allison L Haas, Emily J D’Amico, Catherine R Armbruster, Anna Zemke, Dominique Limoli, Jennifer M Bomberger

**Affiliations:** Department of Microbiology and Molecular Genetics, University of Pittsburgh School of Medicine, Pittsburgh, PA, USA; Division of Infectious Disease, Department of Pediatrics, University of Pittsburgh School of Medicine, Pittsburgh, PA, USA; Department of Biology, Saint Vincent College, Latrobe, PA, USA; Department of Microbiology and Immunology, Carver College of Medicine, University of Iowa, Iowa City, IA, USA; Division of Pulmonary, Allergy, and Critical Care Medicine, Department of Medicine, University of Pittsburgh School of Medicine, Pittsburgh, PA, USA

**Author notes:** Current Address: Department of Microbiology and Immunology, Geisel School of Medicine at Dartmouth, Hanover, New Hampshire, USA.

## Abstract

Mucosa-associated biofilms are associated with many human disease states, but the mechanisms by which the host promotes biofilm biogenesis remain unclear. In chronic respiratory diseases like cystic fibrosis (CF), *Pseudomonas aeruginosa* establishes chronic infection through biofilm formation. *P. aeruginosa* can be attracted to interspecies biofilms through potassium currents emanating from the biofilms. We hypothesized that *P. aeruginosa* could, similarly, sense and respond to the potassium efflux from human airway epithelial cells (AECs) to promote biofilm biogenesis. Using respiratory epithelial co-culture biofilm imaging assays of *P. aeruginosa* grown in association with CF bronchial epithelial cells (CFBE41o^-^), we found that *P. aeruginosa* biofilm biogenesis was increased by potassium efflux from AECs, as examined by potentiating large conductance potassium channel, BK_Ca_ (NS19504) potassium efflux. This phenotype is driven by increased bacterial attachment and increased coalescence of bacteria into aggregates. Conversely, biofilm formation was reduced when AECs were treated with a BK_Ca_ blocker (paxilline). Using an agar-based macroscopic chemotaxis assay, we determined that *P. aeruginosa* chemotaxes toward potassium and screened transposon mutants to discover that disruption of the high-sensitivity potassium transporter, KdpFABC, and the two-component potassium sensing system, KdpDE, reduces *P. aeruginosa* potassium chemotaxis. In respiratory epithelial co-culture biofilm imaging assays, a KdpFABCDE deficient *P. aeruginosa* strain demonstrated reduced biofilm growth in association with AECs while maintaining biofilm formation on abiotic surfaces. Collectively, these data suggest that *P. aeruginosa* biofilm formation can be increased by attracting bacteria to the mucosal surface via a potassium gradient and enhancing coalescence of single bacteria into microcolonies through aberrant AEC potassium efflux sensed through the bacterial KdpFABCDE system. These findings suggest that electrochemical signaling from the host can amplify biofilm biogenesis, a novel host-pathogen interaction, and that potassium flux could be a potential target for therapeutic intervention to prevent chronic bacterial infections in diseases with mucosa-associated biofilms, like CF.

**Author Summary:** Biofilm formation is important for *Pseudomonas aeruginosa* to cause chronic infections on epithelial surfaces during respiratory diseases, like cystic fibrosis (CF). The host factors that promote biofilm formation on host surfaces are not yet fully understood. Potassium signals from biofilms can attract *P. aeruginosa*, but it is unknown if potassium from the human cells can influence *P. aeruginosa* biofilm formation on a host surface. We found that *P. aeruginosa* biofilm formation on human airway cells can be increased by the potassium currents from airway cells, and determined bacterial genes related to potassium uptake and sensing that contribute to biofilm formation on airway cells. These findings suggest that *P. aeruginosa* can respond to host potassium signals by forming increased biofilm and that reducing chronic infections may be accomplished by reducing potassium coming from airway cells or blocking the bacterial proteins responsible for the biofilm enhancement by potassium currents.

## Introduction

Biofilms are communities of bacteria encased in an extracellular matrix of protein, polysaccharides, and extracellular DNA and associated with bacterial tolerance to immune recognition, immune killing, and antibiotic treatment [1, 2]. Biofilms can form on abiotic surfaces like plastic and glass but can also form in association with mucosal surfaces in many diseases, including chronic respiratory diseases such as chronic obstructive pulmonary disease (COPD), cystic fibrosis (CF), and chronic rhinosinusitis [3–6]. While much is understood about biofilm formation on abiotic surfaces, little is known about the host factors regulating biofilm growth in the host. The biofilm growth of some bacteria, like *Pseudomonas aeruginosa*, can be altered by host hormones and cytokines. Specifically, biofilms can be dispersed by human atrial natriuretic peptide through specific transcriptional alterations and interferon-γ can promote adhesion via the OprF sensor [7, 8]. *P. aeruginosa* adhesion can also be influenced by host ions like phosphate altering expression of the surface molecule, PstS. In CF, prior work has shown that the CF airway epithelium supports biofilm growth [9]. Further work demonstrated that iron chelation reduced biofilm growth, suggesting that biofilm formation on the epithelium was partially mediated by iron bioavailability from CF cells [10]. While iron bioavailability in CF is one factor promoting biofilm growth, additional host factors likely regulate biofilm growth on mucosal surfaces. Determining the additional epithelial factors that modulate biofilm biogenesis at mucosal surfaces could improve care of diseases where biofilm growth contributes to the burden of chronic infection.

*P. aeruginosa* is an opportunistic pathogen that causes a range of infections including ventilator associated pneumonia, skin and soft tissue infection, and chronic respiratory infection in CF [11–15]. *P. aeruginosa* establishes chronic infections in CF through biofilm growth in the lungs [5, 16]. About 60% of CF patients will go on to develop chronic infection with *P. aeruginosa* by adolescence or early adulthood [17]. Chronic *P. aeruginosa* infection in CF is associated with poor clinical outcomes, so aggressive treatments are used to try to eradicate infections with courses of antibiotics [18–22]. Recurrent pulmonary exacerbations lead to further antibiotic courses, resulting in the emergence of antibiotic resistant *P. aeruginosa* over time [23]. For all these reasons, chronic respiratory infections with *P. aeruginosa* remain a significant hurdle in the treatment of people with CF [17, 24]. Therefore, understanding the factors that promote chronic infection in the airway is important to CF treatment and can shed light on chronic bacterial infections in other disease states.

CF is a genetic disease resulting from a mutation in the cystic fibrosis transmembrane conductance regulator (CFTR) anion channel [25, 26]. CFTR secretes chloride into the airway surface liquid (ASL) which helps to maintain airway surface liquid hydration [27]. When CFTR is dysfunctional, as in CF, the airway surface liquid becomes dehydrated, collapsing cilia and reducing their ability to beat to clear mucus from the airway. Poor mucociliary clearance allows for debris and bacteria to remain in the lungs, instigating chronic inflammation and infections that result in bronchiectasis, lung function decline, and eventually lung failure [26].

Potassium, an important ion for proper host cellular function, can play an important role in bacterial behavior outside the host, but how host derived potassium regulates bacterial interactions is unknown. Potassium transporters in many species have been shown to regulate virulence factors, stress responses, and biofilm formation [28]. In *P. aeruginosa*, it has been shown that loss of potassium transporters can decrease antibiotic resistance, biofilm formation, and motility [29–31]. Interestingly, potassium currents from biofilms of *Bacillus subtilis* can influence the activity of *B. subtilis* by increasing directional tumbling motility of planktonic *B. subtilis* toward the biofilm [32]. It was also shown that this phenomenon was species independent by demonstrating planktonic *P. aeruginosa* attraction to and incorporation into *B. subtilis* biofilms [32]. The response of *P. aeruginosa* to potassium from *B. subtilis* biofilms led us to hypothesize that host potassium efflux from epithelial cells could similarly attract *P. aeruginosa* and enhance biofilm formation on the epithelial surface. Determining how *P. aeruginosa* responds to potassium from airway epithelial cells could provide new insights into a novel mechanism by which biofilm biogenesis may be altered by the host through electrochemical signaling.

In the current study, we utilized a respiratory epithelial co-culture biofilm live-cell imaging system to investigate if *P. aeruginosa* biofilm biogenesis is regulated by epithelial potassium efflux. Using potassium channel potentiators and inhibitors, we demonstrate that biofilm biogenesis is mediated by host electrochemical signaling, revealing a novel host-pathogen interaction. We further identify a *P. aeruginosa* operon necessary for the potassium-induced biofilm biogenesis on epithelial cells that could serve as a potential target for therapeutic intervention to reduce chronic infections in people with diseases like CF.

## Results

### *P. aeruginosa* biofilm biogenesis is increased by potassium efflux from the respiratory epithelium

Previous work reported *P. aeruginosa* motility toward and incorporation into formed biofilms of *B. subtilis*, an unrelated bacterial species, could be induced by potassium currents from *B. subtilis* biofilms [32]. We sought to determine if potassium efflux from human epithelial cells could induce a similar attraction and biofilm enhancement in *P. aeruginosa*.

To assess the effects of potassium efflux from respiratory epithelial cells, we utilized a respiratory epithelial co-culture biofilm model that we have previously described [9, 33, 34] (Fig 1A). Briefly, airway cells are grown and polarized, then affixed into a chamber where a continuous flow of human cell culture media is perfused. To quantify biofilm biomass, nine randomly selected visual fields are imaged 3-dimensionally at each time point. Bacterial biomass is calculated using the Nikon Elements software algorithm that quantifies volumes of thresholded objects in the images. Images are taken at 1-, 3-, and 6-hours post-inoculation as epithelial barrier integrity is compromised after 6 hours. This model system has been shown to produce biofilms that express the quorum sensing gene *rhlA* and biofilm specific gene *tolA* [9], as well as recapitulate the size and morphology of *P. aeruginosa* aggregates in CF sputum and gene expression of *P. aeruginosa* in CF sputum [35–38].

**Figure 1:**
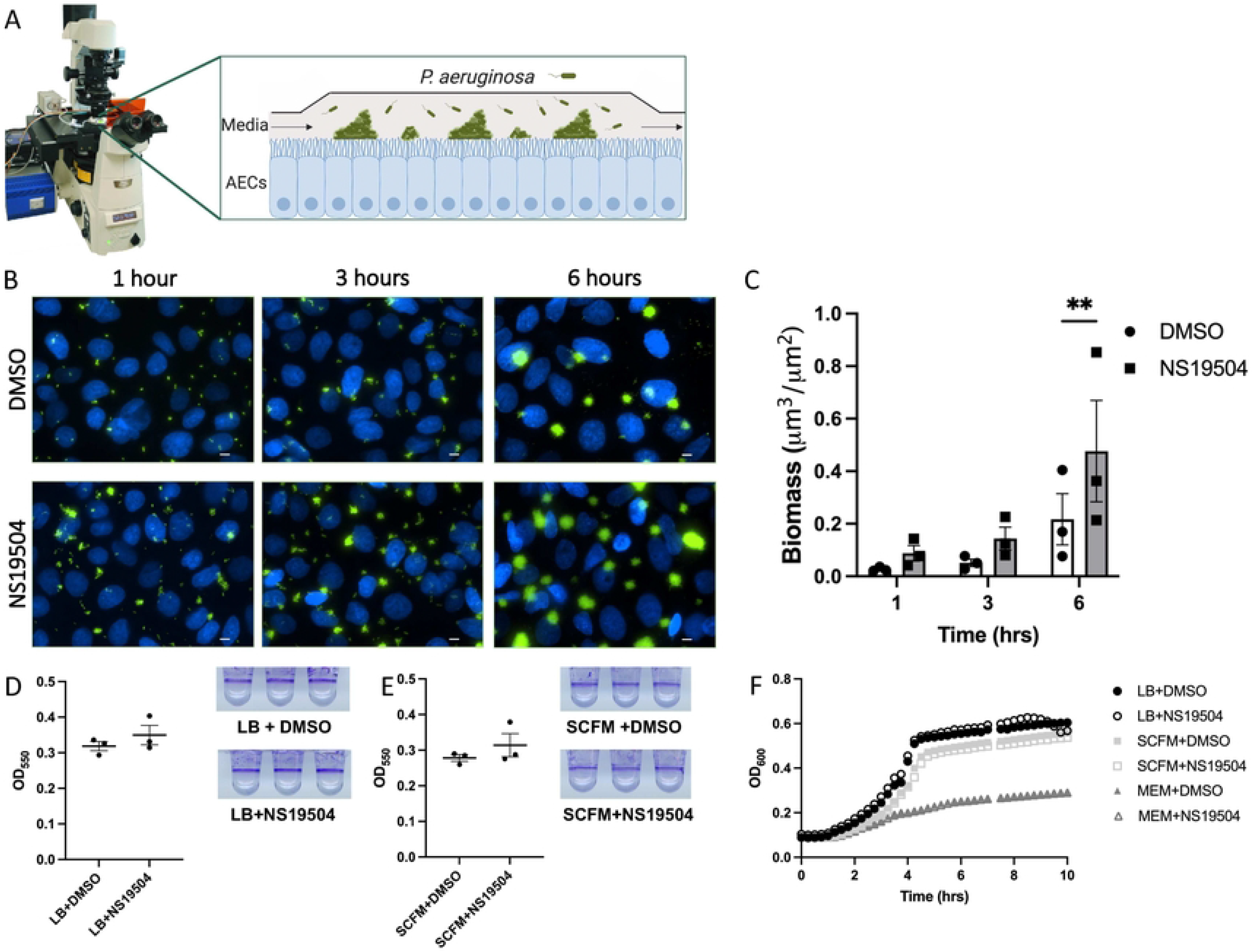
*P. aeruginosa* biofilm formation is enhanced by increased potassium efflux from airway epithelial cells by opening of BK_Ca_ channels. **(A)** Graphical representation of respiratory epithelial co-culture biofilm assay utilized for all imaging studies. **(B)** CFBE41o^-^ cells on glass coverslips were infected with GFP-producing *P. aeruginosa* (green) while stimulated with NS19504 (25 µM), a BK_Ca_ channel potentiator, or 0.05% DMSO in MEM without phenol red and imaged by fluorescent microscopy at 1-, 3-, and 6-hours. CFBE41o^-^ cell nuclei are stained with Hoescht33342 (blue). Scale bar represents 10 μm. **(C)** Biomass (μm^3^/μm^2^) measurements at 1-, 3-, and 6-hours post-inoculation from three independent experiments. Line in bar represents mean value and error bars represent standard error of the mean. Statistical significance was tested by two-way ANOVA with multiple comparisons (* p<0.05). (D) *P. aeruginosa* biofilms grown in 96-well plates in LB and **(E)** SCFM with or without NS19504 measured using crystal violet absorbance at 550 nm. **(F)** Planktonic growth kinetics of *P. aeruginosa* grown in LB Lennox broth (LB), minimal essential media (MEM), and synthetic cystic fibrosis sputum media (SCFM) with or without NS19504.

Multiple potassium channels are expressed on the apical surface of airway epithelial cells. Of these, the large conductance, calcium activated, and voltage-dependent potassium channels (BK_Ca_) have been shown to be important in maintaining airway surface liquid [39]. Several BK_Ca_ channel potentiating compounds exist, have been well-studied and can be used to increase epithelial potassium efflux. In order to increase potassium efflux from the CF bronchial epithelial cell line, CFBE41o-, used in our respiratory epithelial co-culture biofilm assay, we utilized the specific BK_Ca_ channel opener, NS19504 [40–43]. We treated CFBE41o-cells with media containing NS19504 or DMSO, as the vehicle control, throughout the course of a 6-hour biofilm assay. The biomass of *P. aeruginosa* biofilms grown in association with CFBE41o-cells was increased when epithelial potassium efflux was augmented using NS19504, as compared to the vehicle control treated cells (Fig 1B and 1C). After 6 hours of biofilm growth, we observed significantly higher biomass measured when cells were treated with NS19504. To assess if this difference was seen due to the presence of airway cell potassium rather than a direct effect of NS19504 on the bacteria, microtiter dish biofilm assays and planktonic growth curves were evaluated in the presence and absence of NS19504. In the microtiter dish biofilm assay, bacteria form biofilms on wells of a microtiter dish without the presence of airway cells by attaching directly to the plastic wells at the air-liquid interface. Biofilms can then be quantified by staining with crystal violet. In this assay, biofilm growth was not altered by the presence of NS19504 in either LB or synthetic cystic fibrosis sputum (SCFM) media, as seen by the images and measurements of crystal violet staining (Fig 1D and 1E). Moreover, NS19504 did not affect the planktonic growth rate of *P. aeruginosa* in LB, SCFM, or human cell culture media, minimal essential media (MEM), as seen by the almost identical growth curves in the presence or absence of NS19504 (Fig 1F).

To assess if NS19504 was toxic to CFBE41o-cells, cultures differentiated at air-liquid interface (ALI) were treated with NS19504 for 6 hours and transepithelial resistance (TEER) was measured, as a readout for barrier integrity. NS19504 did not alter TEER, as compared with vehicle control treated cells over the 6-hour time course (Fig S1A). We also assessed if NS19504 altered the cytotoxicity of *P. aeruginosa* by measuring TEER and using a 3-(4,5-dimethylthiazol-2-yl)-2,5-diphenyltetrazolium bromide (MTT) based cell viability assay during *P. aeruginosa* infection of our ALI CFBE41o-cells. There was no significant difference between the DMSO or NS19504 treated cells in the MTT assay or in change in TEER at the end the 6-hour infection (Fig S1B and S1C). Taken together, these data suggest that potassium efflux from airway epithelial cells, and not off target effects of NS19504, increases biofilm growth by *P. aeruginosa*.

To determine if the increase in biofilm growth could be replicated by increasing growth media potassium concentrations, in the absence of the respiratory epithelial potassium conductance and potassium gradients, we examined *P. aeruginosa* planktonic and biofilm growth with increasing concentrations of potassium. Using a modified M9 chemically defined media that did not contain potassium, supplemented with potassium chloride, we tested planktonic growth in the setting of different concentrations of potassium. Planktonic growth of *P. aeruginosa* was not altered by differing potassium concentrations, as seen by the similar growth curves except for the 0 mM potassium condition (Fig S2A). In addition, biofilm growth in the microtiter biofilm assay was also not altered by increasing potassium concentrations (Fig S2B). These data suggest that potassium does not directly alter biofilm growth, suggesting that it is the gradient of potassium emanating from airway epithelial cells that increased biofilm growth.

### *P. aeruginosa* attachment is augmented by potassium efflux

Biofilm formation occurs through multiple stages: bacterial attachment, microcolony formation, maturation, and detachment [44, 45]. To better understand which stages of biofilm formation are altered by potassium efflux, deeper analysis of the respiratory epithelial co-culture biofilm assays was conducted. Again, we treated CFBE41o-cells with media containing NS19504 or DMSO, as the vehicle control, throughout the course of a 6-hour infection and images were acquired at 1, 3, and 6 hours after infection and analyzed for bacterial attachment, and the size and number of biofilms. Attachment measurements at 1-hour post-infection demonstrated that there were more bacteria associated with the epithelial cell layer early in the infection with increased potassium efflux from airway cells treated with NS19504, as compared to the DMSO treated (Fig 2A). At the 6-hour time point, there was no significant difference in the number of biofilm aggregates or area of aggregates between when cells were treated with NS19504 versus DMSO treated cells (Fig 2B and 2C). Given that the overall biomass was increased with increased potassium conductance due to NS19504 treatment, and this measurement includes a volumetric measurement, the height of the biofilms must be increased with NS19504 treatment. These data suggest that potassium increases biofilm biogenesis by promoting bacterial attachment, though these current experiments could not assess how potassium may alter microcolony formation.

**Figure 2:**
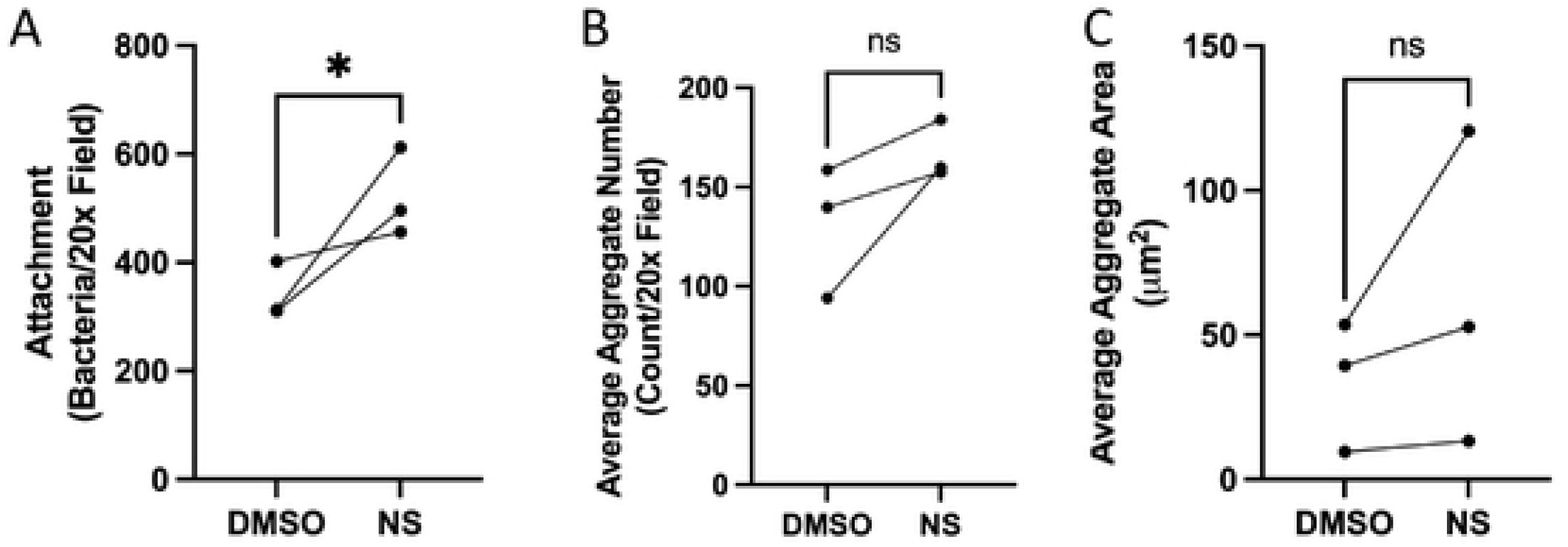
Bacterial attachment, final biofilm size, and final biofilm number are increased by increased potassium efflux. **(A, B, C)** Bacterial attachment at 1 hour and average aggregate area and number at 6 hours grown on CFBE41o^-^ cells in co-culture experiments were measured using Nikon Elements. **(A)** Number of bacteria attached per 20x field for epithelial cells treated with 0.05% DMSO or NS19504 (25 µM) during respiratory epithelial co-culture biofilm experiments at 1 hour. Line represents mean and error bars represent standard error of the mean. **(B)** Average aggregate area per 20x field measured at 6-hour time point for epithelial cells treated with 0.05% DMSO or NS19504 (25 µM) during live-cell co-culture experiments. Line represents mean and error bars represent standard error of the mean. **(C)** Average aggregate number per 20x field measured at 6-hour time point for epithelial cells treated with 0.05% DMSO or NS19504 (25 µM) during live-cell co-culture experiments. Line represents mean and error bars represent standard error of the mean. Line connecting data points indicates data points from same biologic replicate. Statistical significance was tested by unpaired t-test (* p<0.05) for all panels.

### *P. aeruginosa* coalescence into microcolonies is increased by epithelial potassium efflux

We hypothesized that potassium gradients generated by the airway epithelium may also be increasing *P. aeruginosa* motility to form microcolonies, in addition to attachment. We assessed the coalescence of bacteria into microcolonies in association with the respiratory epithelium and if this facet of biofilm growth was impacted by epithelial potassium efflux. We hypothesized that potassium efflux boosts bacterial microcolony formation by increasing bacterial coalescence into microcolonies. In order to test this hypothesis, three different wild-type PAO1 strains expressing different fluorescent proteins, yellow fluorescent protein (YFP), teal fluorescent protein (TFP), and tdTomato, a red fluorescent protein, were used to infect during the co-culture experiments. In the setting of single-color infections, it is impossible to tell if bacteria have come together to form a biofilm or if single bacteria have simply proliferated to grow into the biofilms. By mixing the three different colored strains of PAO1, we were able to determine whether biofilms were formed by multiple individual attached bacteria coming together on the epithelial surface (polyclonal) or by replication of a single clone (monoclonal). In these experiments, we observed that *P. aeruginosa* primarily attached as singleton bacteria at 1 hour, but by 6 hours of growth, a mixture of single color and multicolor biofilms were present (Fig 3A and 3B). In these images, more multicolor biofilm aggregates are present in the group treated with NS19504, as compared to those treated with DMSO. We quantified the proportion of multicolor biofilms at 6 hours in Nikon Elements software by defining a binary which contained all three colors of fluorescence and then using the count objects function to count objects that contained one or more than one fluorescent signal above the threshold value. Biofilms of *P. aeruginosa* in the condition of increased host potassium efflux were more likely to be formed by coalescence of multiple clones (i.e., multicolor biofilms), as compared to cells with basal potassium flux, as seen by the increased proportion of multicolor aggregates noted in the NS19504 treated group (Fig 3C). These findings suggest that epithelial potassium efflux increases biofilm growth by both promoting bacterial association with the respiratory epithelium and increasing microcolony formation by bacterial coalescence on the epithelial surface.

**Figure 3.**
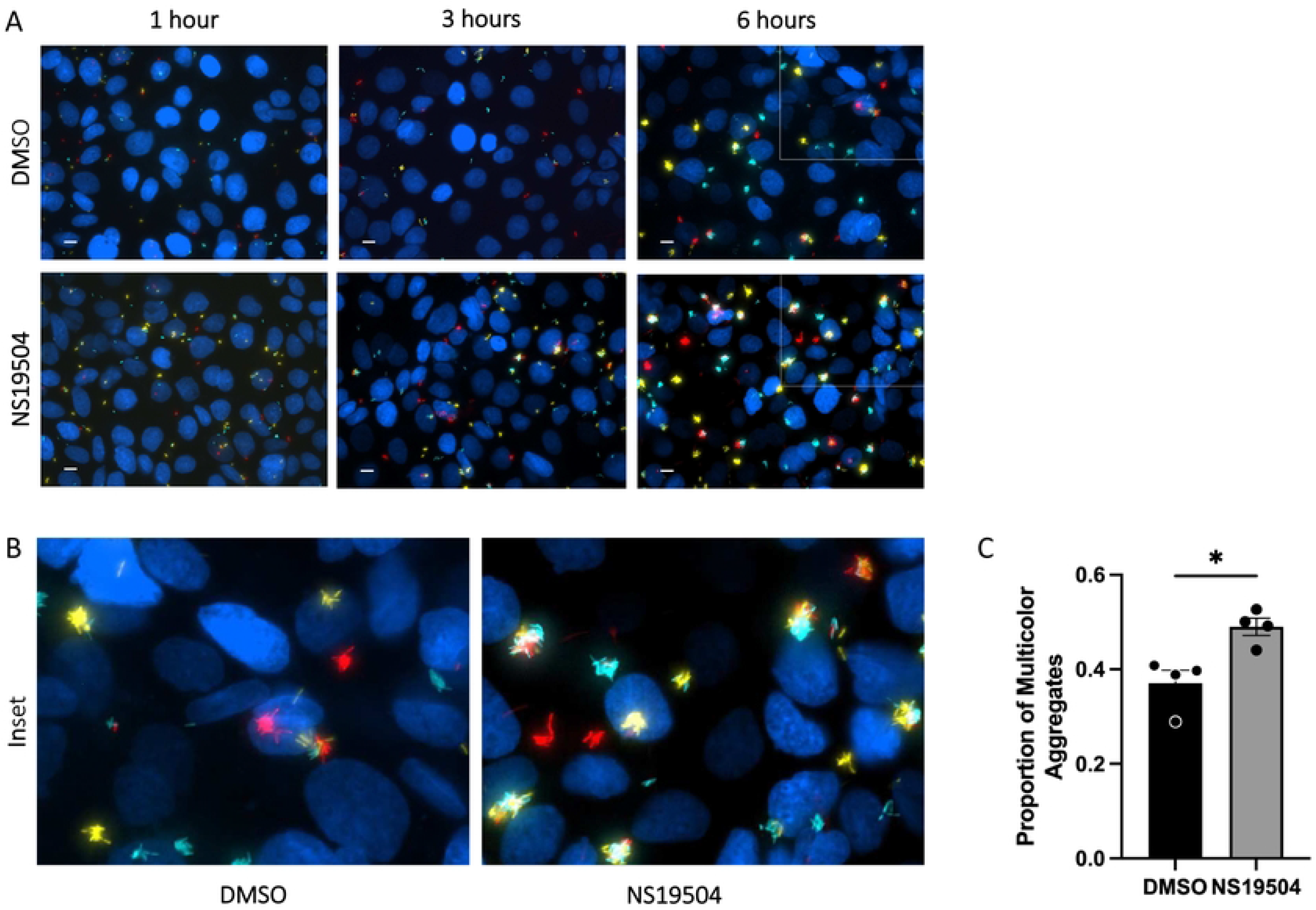
Bacterial coalescence into biofilms is increased by epithelial potassium efflux. **(A and B)** CFBE41o^-^ cells on glass coverslips are infected with a 1:1:1 mixture of TFP-, YFP-, and tdTomato-producing *P. aeruginosa* while treated with NS19504 (25 µM) or 0.05% DMSO in respiratory epithelial co-culture biofilm assay and imaged by fluorescent microscopy at 1-, 3-, and 6-hours post-infection. **(A)** Representative images of bacterial coalescence at 1-, 3-, and 6-hours growth. Scale bar represents 10 μm. White box indicates area of magnification in panel B. **(B)** Magnification of one quarter of panel A images demonstrating multicolor aggregates (polyclonal) compared to single color aggregates (monoclonal). **(C)** Proportion of colocalized bacteria at 6 hours measured with Nikon Elements Software. Bar represents mean with dots representing each of the four biologic replicates and error bars represent standard error of the mean. Statistical significance was determined by unpaired t-test (* p<0.05).

### Biofilm biogenesis is reduced when potassium efflux is decreased

With the finding that potassium efflux increases biofilm formation on airway epithelial cells, we hypothesized that reducing potassium flux would decrease biofilm formation. As with BK_Ca_ channel potentiators, well established, specific BK_Ca_ channel inhibitors exist to block potassium efflux, such as the compound paxilline [46–50]. To test the effects of reduced potassium efflux on biofilm formation, respiratory epithelial co-culture biofilm experiments were conducted with paxilline to inhibit BK_Ca_ channel function. In these studies, biofilm biomass was reduced when epithelial potassium efflux was inhibited by paxilline treatment, as compared to DMSO treated controls (Fig 4A and 4B). Paxilline did not affect *P. aeruginosa* biofilm formation in the absence of airway epithelial cells in LB or SCFM as seen by the similar levels of crystal violet staining in the microtiter dish biofilm assay in both the paxilline and DMSO treated bacteria in both media (Fig 4C). Additionally, planktonic growth in any of the three growth media (LB, SCFM, and MEM) tested was not altered by paxilline, as seen by the similar growth kinetics for the paxilline and DMSO treated bacteria (Fig 4D). Paxilline also did not reduce epithelial cell barrier integrity in the absence of bacterial infection as seen by the increasing TEER over the course of 6-hour treatment (Fig S3A). Additionally, with paxilline treatment during bacterial infection, there was no change in cell viability, as measured by MTT assay, or difference in TEER during infection when paxilline treated cells were compared to the DMSO treated (Fig S3B ad S3C). As expected, based on the finding that potassium efflux increased bacterial attachment, reduced potassium flux from paxilline treatment resulted in decreased bacterial attachment, but did not decrease final biofilm number or area (Figure S4A-C). These data indicate that biofilm biogenesis in association with airway epithelial cells (AECs) can be reduced by reducing the potassium flux from the airway cells.

**Figure 4:**
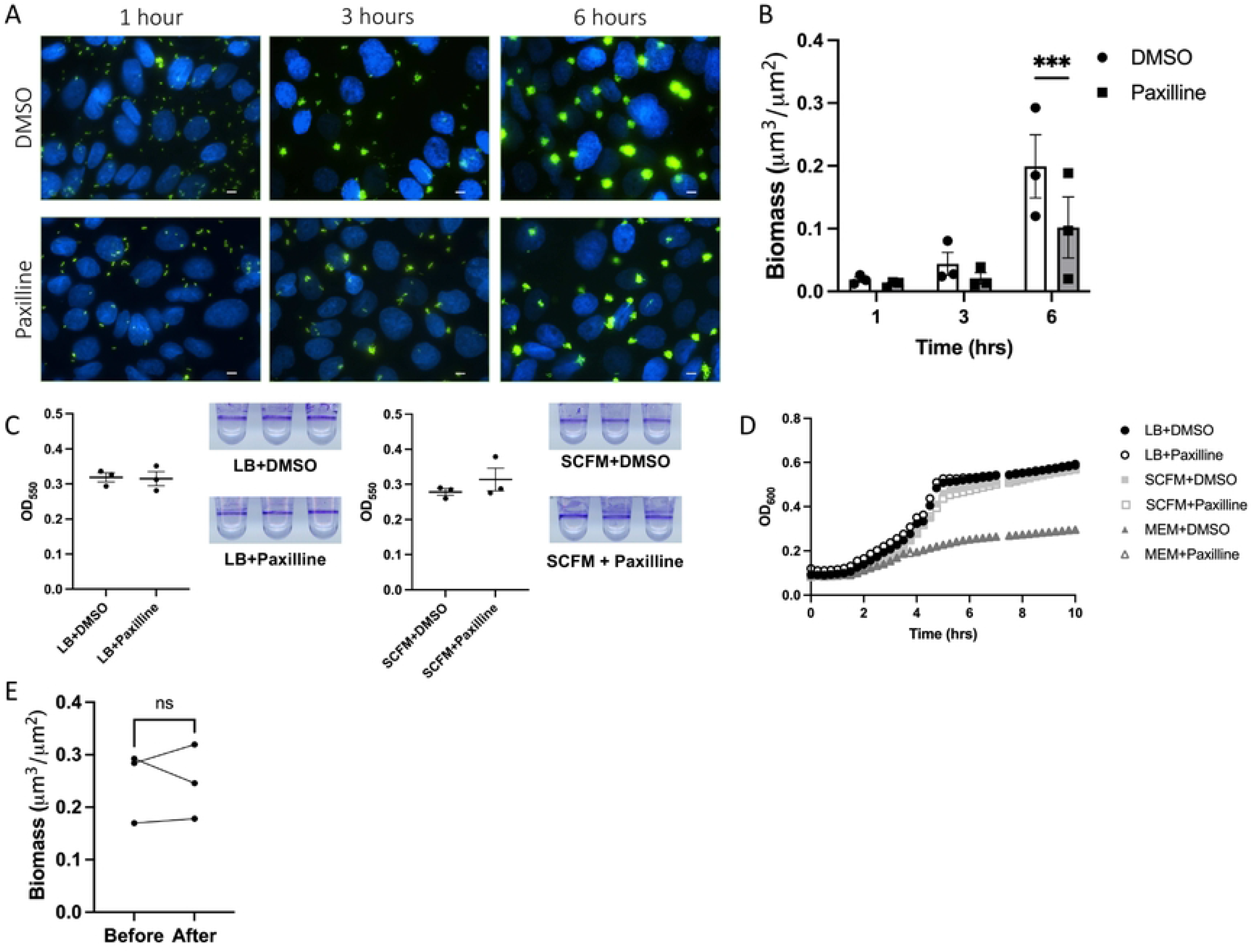
*P. aeruginosa* biofilm formation is decreased by reducing potassium efflux from airway epithelial cells through inhibition of BK_Ca_ channels. **(A)** CFBE41o^-^ cells on glass coverslips were infected with GFP-producing *P. aeruginosa* (green) while treated with paxilline (10 µM), a BK_Ca_ channel inhibitor, or DMSO at 0.05% in MEM without phenol red and imaged by fluorescent microscopy at 1, 3, and 6 hours. CFBE41o^-^ cell nuclei are stained with Hoescht33342 (blue). Scale bar represents 10 μm. **(B)** Biomass (μm^3^/μm^2^) measurements (measured with Nikon Elements Software) at 1-, 3-, and 6-hours post-inoculation from three independent experiments. Line in bar represents mean value and error bars represent standard error of the mean. Statistical significance was tested by 2way ANOVA with multiple comparisons (*** p<0.001). (C) *P. aeruginosa* biofilms grown in 96-well plates in LB and SCFM with or without paxilline measured using crystal violet absorbance at 550 nm. **(D)** Planktonic growth kinetics of *P. aeruginosa* grown in LB Lennox broth (LB), minimal essential media (MEM), and synthetic cystic fibrosis sputum media (SCFM) with or without paxilline. **(E)** Biomass (μm^3^/μm^2^) measurements before and after paxilline treatment. After biofilms were formed and measured on DMSO treated cells for 6 hours, paxilline containing media was introduced for 30 minutes and biofilms were measured. Line represents mean and error bars represent standard error of the mean. Line connecting data points indicates data points from same biologic replicate.

Given the effect of paxilline on reducing biofilms, we examined if paxilline could function as a dispersal agent for established biofilms. In order to assess this, biofilms were grown in the presence of MEM with DMSO for 6 hours and imaged. The media was then changed to MEM containing paxilline and perfused for 30 minutes prior to re-imaging. We have previously published the characterization of this model for studying biofilm dispersal [51]. Biomass was unchanged after the 30 minutes of paxilline treatment suggesting that loss of potassium efflux from the respiratory epithelium does not function as a dispersal signal (Fig 4E). This data suggests that epithelial cell potassium gradients are important in the establishment of biofilms, but do not affect their dispersal.

### *Kdp* potassium sensing and uptake genes are necessary for chemotaxis toward potassium

After demonstrating that epithelial potassium efflux enhances biofilm formation by *P. aeruginosa*, we next identified the bacterial mechanisms involved in sensing and responding to potassium. Based on the previous work [32] and our findings in this study, we hypothesized that potassium was leading to increased bacterial motility toward the potassium current. To test this hypothesis, we utilized a previously described macroscopic twitching chemotaxis assay with potassium chloride as the chemoattractant [52–54]. We observed twitching chemotaxis toward the potassium gradient with wild-type PAO1, while a *ΔpilA* strain of PAO1, deficient in twitching motility, did not chemotax toward potassium. This was observed as greater expansion of bacteria toward the potassium gradient on the left side of the growth ring in the wild-type PAO1 and lack of this expansion in any direction in the *ΔpilA* strain (Figure 5A). Additionally, this *P. aeruginosa* showed an increased directional motility in their chemotaxis toward the potassium chloride gradient, as compared to a control water treatment (Figure 5A, striped bar).

**Figure 5.**
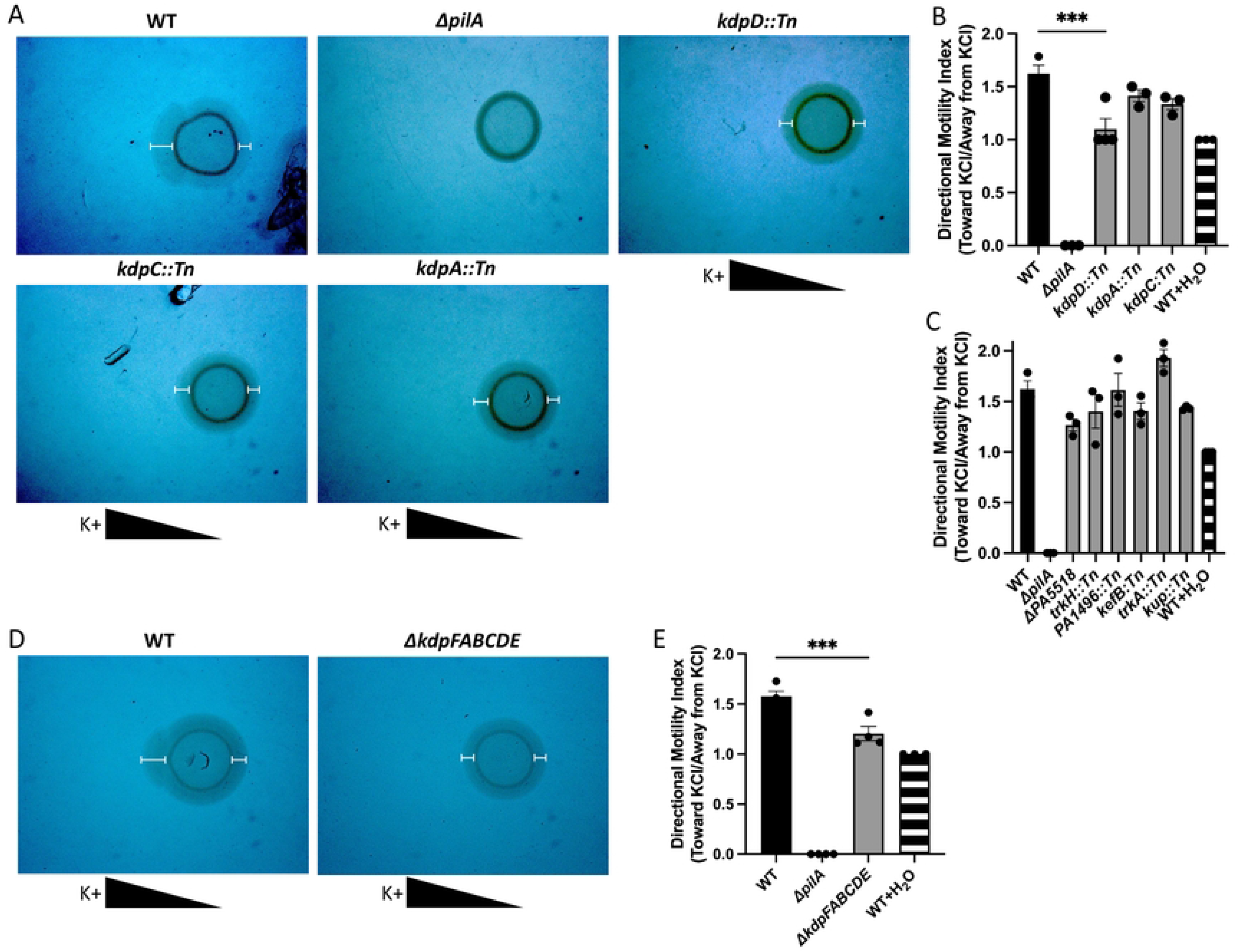
*P. aeruginosa* chemotaxis toward a potassium gradient is reduced by deletion of *kdp* genes. **(A)** Stereomicroscopy images of macroscopic twitching chemotaxis assay demonstrated WT MPAO1 *P. aeruginosa* has directional motility towards potassium chloride (KCl) gradient. Transposon mutants in the *kdp* genes (*kdpD, kdpC, kdpA*) have reduced directional chemotaxis toward potassium. *ΔpilA* mutant served as negative control for twitching motility. Triangles below images represent the KCl gradient and white I-lines on images were applied to enhance visualization of twitching motility. **(B)** Directional motility index measured by the ratio of motility toward the potassium gradient over the motility away from the potassium gradient for the *kdp* gene transposon mutants tested. WT with water served as negative control for directional chemotaxis. Statistical significance tested by ANOVA (** p<0.01). **(C)** Directional motility index measured for other potassium related genes in the transposon mutant library and a markerless, in-frame deletion of *PA5518*. **(D)** Stereomicroscopy images of macroscopic twitching chemotaxis assay demonstrated WT PAO1 *P. aeruginosa* has directional motility toward potassium gradient while a strain with a markerless, in-frame deletion of *kdpFABCDE* has reduced directional motility. Triangles below images represent the KCl gradient and I-lines on images were applied to enhance visualization of twitching motility. **(E)** Directional motility index of WT PAO1 compared to *ΔkdpFABCDE* strain. Statistical significance was tested by unpaired t-test (** p<0.01)

Having shown that *P. aeruginosa* chemotaxes toward potassium and that this chemotaxis is pilus mediated, we interrogated the *P. aeruginosa* genome for genes annotated to be associated with potassium transport [55]. The search yielded 12 genes of interest, 6 of which were part of the *kdp* gene cluster. Available transposon mutant strains with insertions in the identified potassium transport genes from an MPAO1 transposon mutant library and a POA1 *P. aeruginosa* strain with a markerless, in-frame deletion of PA5518 were screened for chemotaxis toward potassium using the macroscopic twitching chemotaxis assay [56, 57]. We observed that mutants in the *kdp* gene cluster, particularly the *kdpD* transposon mutant, had reduced chemotaxis toward potassium, as seen by decreased expansion of these mutants toward the potassium gradient corresponding to a lower directional motility index, while the other mutants screened did not have a reduction in chemotaxis (Fig 5A, 5B, and 5C). *kdpFABC* encodes a high affinity potassium uptake channel and *kdpDE* encodes an intracellular potassium sensor kinase, KdpD, and a response regulator that is phosphorylated by KdpD, KdpE [58–60]. Little is known about this two-component system in *P. aeruginosa*, but in other species, KdpE has been shown to regulate virulence factors and growth [59, 61, 62].

With the evidence that the *kdp* genes are involved in *P. aeruginosa* chemotaxis toward potassium, we generated a PAO1 strain with a markerless, in frame deletion of all *kdp* genes (Δ*kdpFABCDE)* using a previously described mutagenesis protocol [63] and confirmed this deletion by whole genome sequencing. We assessed this mutant in the chemotaxis assay and observed a significant reduction in chemotaxis toward potassium, as compared to the wild-type PAO1 control (Fig 5D and 5E). With the knowledge that *P. aeruginosa* chemotaxes toward potassium and that deletion of the *kdp* operon reduces potassium chemotaxis, we next examined biofilm biogenesis in our respiratory epithelial co-culture biofilm system to determine if the *kdp* operon may be involved in the potassium-induced biofilm formation in association with the CF airway epithelium.

### *Kdp* genes regulate biofilm formation on epithelial cells

Before evaluating the role of the *kdp* genes in our respiratory epithelial co-culture biofilm assay, we investigated if the Δ*kdpFABCDE* strain was deficient in planktonic or abiotic biofilm growth as compared wild-type PAO1. Planktonic growth curves showed similar growth kinetics between the Δ*kdpFABCDE* and its wild-type PAO1 parental strain in LB broth, SCFM, and MEM (Fig. S5A). Biofilm growth in the microtiter dish biofilm assay (i.e. abiotic biofilm growth) was similar in LB and SCFM (Fig. S5B). To ensure that the deletion did not alter general twitching motility, a twitching motility assay was performed. The D*kdpFABCDE* was not deficient in twitching motility when compared to wild-type PAO1, as seen by the similar twitching radius in the WT and D*kdpFABCDE* (Fig. S5C).

Finally, we tested the Δ*kdpFABCDE* strain in our respiratory epithelial co-culture biofilm assay. We observed that the Δ*kdpFABCDE* mutant *P. aeruginosa* strain was deficient in biofilm biogenesis in association with epithelial cells, as evidence by the reduced GFP signal in imaging and reduced biofilm biomass quantified (Fig 6A and 6B). Additionally, we tested the Δ*kdpFABCDE* mutant in the presence of NS19504 which did not rescue the phenotype, suggesting that the Kdp operon is necessary to sense airway epithelial potassium gradients and respond by inducing biofilm growth (Fig 6A and 6B). The Δ*kdpFABCDE* mutant had reduced attachment to the airway epithelium as seen by the reduced bacterial counts at 1-hour post-infection, as well as reduced aggregate area and aggregate number at 6-hours of growth (Fig 6C-6E). These findings demonstrate that the *kdp* genes, involved in potassium sensing and uptake, mediate biofilm biogenesis in association with the airway epithelium and the induced biofilm response to enhanced potassium efflux produced by treatment with NS19504.

**Figure 6.**
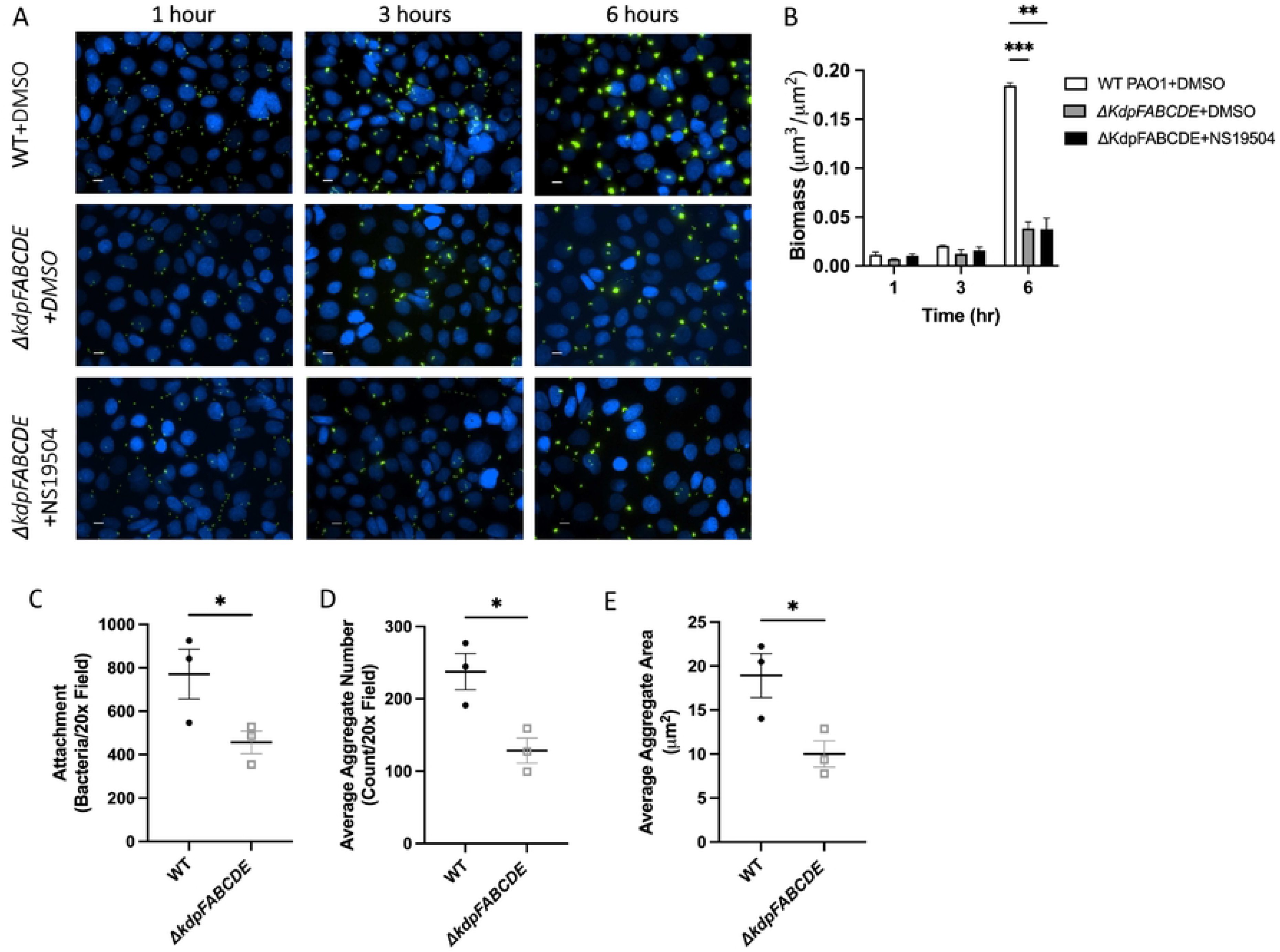
KdpFABCDE deficient *P. aeruginosa* has reduced biofilm formation on AECs. **(A)** CFBE41o^-^ cells on glass coverslips were infected with wild type (WT) PAO1 or *ΔkdpFABCDE* GFP-producing *P. aeruginosa*, grown with continuous flow of MEM with either DMSO or NS19504 and imaged by fluorescent microscopy at 1-, 3-, and 6-hours. CFBE41o^-^ cell nuclei are stained with Hoescht33342 (blue). Scale bar represents 10 μm. **(B)** Biomass (μm^3^/μm^2^) (measured with Nikon Elements) at 1-, 3-, and 6-hours post-inoculation from four independent experiments. Statistical significance was tested by 2-way ANOVA with multiple comparisons (** p<0.01, *** p<0.001). **(C-E)** Bacterial attachment at 1 hour and average aggregate area and number at 6 hours grown on CFBE41o^-^ cells in live-cell co-culture experiments were measured using Nikon Elements. **(C)** Number of bacteria attached per 20x field with WT or *ΔkdpFABCDE* GFP-producing *P. aeruginosa* during live-cell co-culture experiments at 1 hour. Line represents mean and error bars represent standard error of the mean. **(D)** Average aggregate area per 20x field measured at 6-hour time point when infected WT or *ΔkdpFABCDE* GFP-producing *P. aeruginosa*. Line represents mean and error bars represent standard error of the mean. **(E)** Average aggregate number per 20x field measured at 6-hour time point with WT or *ΔkdpFABCDE* GFP-producing *P. aeruginosa*. Line represents mean of four biologic replicates and error bars represent standard error of the mean. Statistical significance was tested by unpaired t-test (* p<0.05).

## Discussion

In the current study, we report that potassium efflux from the respiratory epithelium promotes biofilm biogenesis by enhancing *P. aeruginosa* bacterial attachment and coalescence into microcolonies. Furthermore, we demonstrated that the *P. aeruginosa kdp* gene cluster, a putative potassium uptake system and its associated potassium sensor and regulator, were necessary for biofilm growth associated with the airway epithelium. Our findings suggest a novel aspect of host-pathogen interactions whereby bacterial biofilm biogenesis is increased by the electrochemical gradient at the mucosal surface. Additionally, our study purports possible new therapeutic targets for impeding chronic bacterial infections in diseases with mucosa-associated biofilms by targeting epithelial potassium efflux.

The data in the current study shows a link between *P. aeruginosa* biofilm biogenesis and potassium efflux from host cells. Our study builds on previous work demonstrating that *P. aeruginosa* motility can be altered by *B. subtilis* biofilms through an electrochemical signal, specifically potassium current [32]. Similarly, the data presented here indicates that potassium gradients generated from human respiratory cells can influence *P. aeruginosa* behavior to enhance biofilm biogenesis. And based on the present work, the potassium signal from AECs is sensed over the background potassium concentration of cell culture media, a concentration similar to physiologic extracellular levels, suggesting further *in vivo* relevance. Our findings suggest that the importance of potassium is likely mediated by increasing attachment of *P. aeruginosa* possibly through the previously seen increase in motility resulting from potassium gradients [32, 64]. We posit that the gradient of potassium coming from airway epithelial cells enhances *P. aeruginosa* motility in the direction of the potassium gradient, providing rapid movement toward a surface for attachment. This is further supported by the fact that fixed levels of increased potassium do not alter bacterial growth or biofilm formation without the gradient from airway cells. As the first step in biofilm formation, bacterial attachment, and the enhancement thereof, is likely to drive increased biofilm formation. Potassium also serves many important functions during biofilm growth and development. It has been shown that potassium helps coordinate biofilm growth by coordinating metabolic activities between internal and peripheral organisms in the biofilm through membrane potential [65]. It is possible that the locally higher levels of potassium at the cell surface in the setting of increased potassium efflux with NS19504 improves this metabolic coordination.

Biofilms are found in close association with airway epithelial cells in CF, but they are also found in smaller aggregates associated with the mucus layer. While mucus-bound biofilms and bacteria on the apical side of the mucus are unlikely to see the potassium gradient generated by airway cells, SCFM, which contains 15.8 mM potassium, was created based on chromatographic analysis of CF sputum, suggesting that potassium is much higher in CF sputum than in other extracellular compartments [66]. As noted above, this increased potassium may help mucus-bound biofilms coordinate metabolic activities [65]. When mucus-bound biofilms disperse, the potassium gradient coming from the underlying airway cells may also draw the now planktonic *P. aeruginosa* to the airway cells by increasing their motility in the direction of the epithelium [32, 64].

Potassium transporter activity regulates virulence factor production, stress responses, and biofilm formation in pathogens such as *Staphylococcus aureus* and *Mycobacterium tuberculosis* [28, 67–74]. In *P. aeruginosa*, the loss of the potassium transporter, TrkA, can decrease antibiotic resistance, biofilm formation, and motility [29–31]. Additionally, in a uropathogenic *Escherichia coli* strain, potassium can induce TIR-containing protein C, a protein that modulates host innate immune responses [75]. Despite these roles for potassium in bacterial behaviors, enhancement of *P. aeruginosa* biofilm biogenesis by potassium efflux from a host epithelium is a novel host-pathogen interaction that should be explored in other species that cause chronic infections.

In our transposon mutant screen, we found that mutant strains with insertions in the *kdp* operon had impaired chemotaxis toward a potassium gradient. We showed that the *kdp* operon genes are necessary for biofilm formation in association with AECs. While little is known about the *kdp* operon in *P. aeruginosa*, the KdpDE two-component system is conserved in many bacterial species [76]. In several bacterial species, the KdpDE two-component system has been shown to upregulate virulence factors like metalloproteases, suggesting that in low potassium environments virulence may be increased to increase nutrient availability [62, 67, 77, 78]. The KdpDE two-component system is also important in bacterial stress responses, another indication of the importance of potassium in host-pathogen interactions [79–81]. In *P. aeruginosa, kdpD* has been associated with enhanced biofilm formation on endotracheal tubes used for mechanical ventilation [82]. Small molecules targeting the KdpFABC potassium transporter or the KdpDE two component system could be utilized as adjunctive therapy during antibiotic therapy, particularly when *P. aeruginosa* is first identified in the airway, to reduce biofilm formation, improve antibiotic activity, and thus reduce chronic infection.

In addition to targeting bacterial proteins involved in potassium uptake and homeostasis, therapeutics could target host potassium to interfere with the enhancement of biofilm biogenesis thereby reducing chronic infection. While antibiotic therapy has been used extensively to reduce the bacterial burden in diseases with chronic infections, like chronic rhinosinusitis and CF, success rates of bacterial clearance are variable [83–85]. Therapies that can inhibit biofilm formation have the potential to reduce chronic infection, antibiotic use, and antibiotic resistance development and improve eradication efforts. The current work and prior work with *B. subtilis* suggest that potassium may drive bacterial movement toward the source of potassium [32] and enhance biofilm formation, therefore altering the potassium environment may impede biofilm formation in the airway.

Altering the potassium environment in the airway could be accomplished in several ways. First, an alternative source of potassium that disrupts the potassium gradient emanating from the respiratory epithelium could deter *P. aeruginosa* from associating with the epithelium to establish biofilms. Inhaled potassium salt solution could provide such an alternative potassium source. Alternatively, based on the findings of this study, a pharmacologic agent that reduces potassium flux from the airway epithelium could interfere with *P. aeruginosa* biofilm biogenesis in the airway. In healthy airways, potassium efflux through apical calcium-activated voltage-dependent potassium channels, BK_Ca_ channels, helps to maintain ASL hydration [39, 86]. But in CF, where there is reduced sodium and chloride, potassium plays an increased role in maintaining the minimal ASL on CF epithelial cells [87]. Mathematical modeling shows a baseline increase in potassium efflux from airway epithelial cells through BK_Ca_ channels in the setting of CFTR dysfunction [87]. These studies suggest that potassium blockade may have even greater role in CF due alterations of the electrochemical gradient from CFTR dysfunction. While studies have shown that reducing BK_Ca_ channel potassium flux could further exacerbate mucociliary dysfunction [39, 87, 88], more recent work has shown that deletion of the gene for a different potassium channel, KCa3.1, improves mucociliary clearance in a mouse model of muco-obstructive lung diseases like CF [89], potentially indicating potassium blockers are a viable therapeutic option. The effects of a pharmacologic agent to block potassium efflux on mucociliary clearance would have to be monitored closely, but these agents are promising as adjunctive agents.

Taken together, our findings suggest a mechanism by which *P. aeruginosa* biofilm formation is increased by the electrochemical environment at mucosal epithelial surfaces. Future studies may build on the current findings by studying electrochemical host-pathogen interactions at other mucosal sites and with other bacterial species, and developing and testing bacterial KdpFABC or KdpDE system inhibitors, human potassium channel inhibitors, or inhaled potassium to inhibit mucosa-associated bacterial biofilm formation in many disease states.

## Materials and Methods

### Bacterial strains and cell lines

All strains utilized for this manuscript are documented in Table S1. For all experiments not requiring imaging, the *P. aeruginosa* strain PAO1 was utilized. For imaging experiments involving PAO1, the *P. aeruginosa* strain PAO1 constitutively expressing green fluorescent protein (GFP) was used as previously described [9, 10, 34]. *P. aeruginosa* PAO1 strains constitutively expressing teal fluorescent protein (TFP), yellow fluorescent protein (YFP), and TdTomato were created as previously described [90] with vectors gifted to our lab by Dr. Boo Shan Tseng Lab. MPAO1 transposon mutants for chemotaxis screen were purchased from Manoil Lab transposon mutant library [56, 57]. A markerless, inframe deletion mutant of *ΔpilA* was kindly gifted by Dr. Matthew Parsek and has been published previously [91]. A markerless, inframe deletion of the *kdpFABCDE* genes and *PA5518* in the PAO1 strain was created using a precision mutagenesis method previously described [63] with primers listed in Table S2. Overnight cultures of bacterial strains were grown in 5 mL of LB Lennox broth (Sigma) at 37°C shaking at 300 RPM.

CFBE41o-cells were cultured in media containing penicillin and streptomycin, except during experiments. The identity and purity of the CFBE41o-cells were verified by short tandem repeat profiling (University of Arizona Genetics Core). The immortalized human CF bronchial epithelial cell line CFBE41o-, isolated from a ΔF508/ΔF508 patient, were seeded on glass coverslips, and grown for 7-8 days prior to live cell biofilm co-culture experiments, as described previously [9, 33, 51]. CFBE41o-cells were cultured on Transwell filters for 7-10 days prior to monitoring for BK_Ca_ channel modulator, NS19504 (opener) and paxilline (inhibitor), toxicity, as described previously [9, 33, 34, 92].

### Reagents

Potassium channel modulating reagents, NS19504 (Cat No. 5276) and paxilline (Cat No. 2006), were purchased from Tocris Biosciences (Minneapolis, MN). Lyophilized powder of NS19504 and paxilline was dissolved in 100% cell culture grade DMSO at stock concentrations and frozen at −20° C for up to 2 months for later dilution in MEM without phenol red. LB Lennox broth was purchased from Sigma Life Sciences. Synthetic cystic fibrosis sputum media was (SCFM) made as previously described [66].

### Respiratory epithelial co-culture biofilm experiments

Live-cell imaging was utilized to image biofilm growth on CFBE41o-cells, as previously described with slight modifications [9, 10]. Briefly, we used a FCS2 closed-system, live-cell imaging chamber (Bioptech, Butler PA). Glass coverslips with CFBE41o-cells grown for 7 days were affixed into the chamber with continuous flow of Minimal Essential Media (MEM) (GIBCO) without phenol red supplemented with 2 mM L-glutamine (GIBCO). In DMSO chambers, MEM without phenol red was supplemented with 0.05% DMSO and for NS19504 and paxilline conditions MEM without phenol red supplemented with final concentration of 25 μM for NS19504 or 10 μM paxilline in DMSO resulting in a final concentration of DMSO in the NS19504 and paxilline groups of 0.05%. An overnight culture of PAO1 or *ΔkdpFABCDE* PAO1 constitutively expressing GFP was diluted to OD_600_ of 0.1 and inoculated into the chambers with flow paused. After 1 hour of still incubation, flow of media was resumed at 20 mL/hour for 15 minutes prior to taking 1-hour images. Images were taken at 1-, 3-, and 6-hour time points with the 1-hour time point representing the attachment phase. Images were taken at 20x magnification for measurements and representative 40x magnification images were taken as well. Biomass volume was measured using Nikon Elements software. Nine visual fields were randomly selected and measured at each time point for each experiment, and the average of these nine visual fields was considered one biologic replicate. Two-way ANOVA with Sidak’s multiple comparisons test was conducted to determine statistical significance in biomass. For experiments with the GFP expressing *ΔkdpFABCDE* PAO1, wild-type PAO1 served as the control and cells were treated with MEM without phenol red and DMSO or NS19504.

For measurements of attachment, final aggregate size, and final aggregate number, these experiments were further analyzed using Nikon elements to measure number of bacteria and aggregates as well as the mean aggregate area. Similar to biomass measurements, nine visual fields were randomly selected and measured at each time point for each experiment, and the mean number of bacteria, number of aggregates, and aggregate area for a given day are reported as one biologic replicate. Unpaired t-test was used to determine statistical significance.

### Planktonic growth assay

Overnight cultures of PAO1 and *ΔkdpFABCDE* were grown in LB broth then washed twice with phosphate buffered saline (GIBCO). Washed cultures were diluted 1:100 in LB, SCFM [66], or MEM without phenol red with 25 μM for NS19504 in DMSO, 10 μM paxilline in DMSO, or 0.05% DMSO as control to determine planktonic bacterial growth in the presence of these chemical modulators. Bacterial dilutions were then plated into a 96-well plate, and the plate was placed in 37°C SpectraMax M2 plate reader. OD_600_ readings were taken from each well every 15 minutes with shaking for 30 seconds before each reading for 10 hours. Growth curves were visualized and analyzed using GraphPad. Three technical replicates were completed for each experiment and the experiment was repeated three times.

### Microtiter dish biofilm assay

Overnight cultures of PAO1 and *ΔkdpFABCDE* were grown in LB broth then washed twice with phosphate buffered saline (GIBCO). After resuspension, the optical density was measured by spectrophotometer and cultures were normalized to equivalent OD_600_. Normalized cultures were diluted 1:30 in LB and SCFM with or without 25 μM for NS19504 in DMSO, 10 μM paxilline in DMSO, or 0.05% DMSO in 96-well vinyl U-bottom plates (Corning). Plates were incubated in a resealable bag with a moistened paper towel to maintain humidity and incubated 37°C for 6 hours. After 6 hours of incubation, media was removed from wells, and wells were washed in deionized (DI) water twice. Plates were dried upside down for 15 minutes, then stained with 41% crystal violet in 12% ethanol in DI water. Crystal violet solution was aspirated, and wells were washed three times with deionized water and allowed to dry overnight. After overnight drying, wells were cut out of the plate. To quantify biofilm formation, 30% acetic acid was added to the wells to dissolve crystal violet from the biofilms and the acetic acid OD_550_ was measured using SpectraMax M2 plate reader. Three technical replicate wells were done for each experiment, and the experiment was repeated three times.

### Cytoxicity assay

Transwell filters seeded with CFBE41o-cells were grown for 7-10 days at air-liquid interface. On the day of the experiment, MEM without phenol red with 25 μM for NS19504, 10 μM paxilline in DMSO, or 0.05% DMSO was placed on the apical side of the Transwell filters. Transepithelial resistance (TEER) was measured immediately after media was added to the apical surface, then again at 1, 3, and 6 hours after media was applied using an epithelial volt/ohm meter (World Precision Instruments). In experiments where cells were infected with PAO1, the same experiments were completed with infection with PAO1 at an MOI of 25 as previously described [33]. At the end of 6-hour period, CyQUANT MTT Cell Viability Assay (Invitrogen) was completed on all wells per manufacturer instructions to determine relative cell viability compared to DMSO treated.

### Bacterial coalescence in biofilm co-culture

Live-cell imagining experiments examining the influence of potassium efflux on bacterial coalescence were conducted as above with slight modification. The bacteria used to infect CFBE41o-coverslips was a 1:1:1 mixture of *P. aeruginosa* strains constitutively expressing teal fluorescent protein, yellow fluorescent protein, and tdTomato (Table S1). Chambers were treated with MEM without phenol red with 25 μM NS19504 or 0.05% DMSO as control. Chambers were imaged at 1, 3, and 6 hours. The number of biofilms containing one or more than one strain of *P. aeruginosa* was calculated using Nikon Elements software. Nikon Elements uses Boolean logic based on input threshold values for each fluorophore to determine which biofilms contain signal from more than fluorophore. We reported these results as the percent of total aggregates that had more than one strain of *P. aeruginosa.* The proportion of aggregates with 2 or more colors of *P. aeruginosa* in the aggregate out of the total number of aggregates was measured in nine visual fields at each time point. These nine measurements were averaged and were considered one biologic replicate. The experiment was repeated four times and the mean proportion of multicolor aggregates is reported. Unpaired t-test was used to determine statistical significance.

### Potassium macroscopic chemotaxis assay

A macroscopic twitching chemotaxis assay was adapted from the assay previously described with slight modifications [52–54]. Potassium chloride at a concentration of 300 mM was spotted on buffered minimal agar 4 mm from the site where *P. aeruginosa* strains were to be spotted. A *ΔpilA* strain of *P. aeruginosa* PAO1 served as a twitching motility negative control and sterile deionized water served as a negative chemotaxis control. For transposon mutant screening, MPAO1 parental strain from the Manoil Lab library was utilized as the WT control [56, 57]. For experiments assessing chemotaxis of *ΔkdpFABCDE* PAO1 strain, PAO1 Parsek wild-type strain was used as WT bacterial control.

### Twitching motility assay

Overnight cultures of WT PAO1 and *ΔkdpFABCDE* PAO1 strain were grown in M9 broth media. Cultures were then inoculated through M9 media supplemented with 1.5% agar to the plastic surface of the petri plate using a sterile 10 μL pipette tip. Cultures were allowed to grow for 48 hours. After growth, agar was carefully removed from the petri dish as to not disrupt the bacteria under the agar. Bacterial growth on the petri plate was stained with 10% crystal violet, then washed three times in deionized water. Experiments were done in duplicate and repeated four separate times for four biologic replicates. Plates were imaged using stereomicroscope and twitching radius was measured using Nikon Elements software.

## Acknowledgements

We thank Dr. Boo Shan Tseng for the contribution of the fluorescent protein vectors, and Dr. Matthew Parsek for the *ΔpilA* PAO1.

